# Uveal Melanoma: a miR-16 disease?

**DOI:** 10.1101/2021.11.30.470499

**Authors:** Anaïs M. Quéméner, Laura Bachelot, Marc Aubry, Stéphane Avner, Delphine Leclerc, Gilles Salbert, Florian Cabillic, Didier Decaudin, Bernard Mari, Frédéric Mouriaux, Marie-Dominique Galibert, David Gilot

## Abstract

Uveal melanoma (UM), the most common primary intraocular tumor in adults, has been extensively characterized by omics technologies during the last 5 years. Despite the discovery of gene signatures, the molecular actors driving the cancer aggressiveness are not fully understood and UM is still associated to a dismal overall survival at metastatic stage. Here, we showed that microRNA-16 (miR-16) is involved in uveal melanoma by an unexpected mechanism. By defining the miR-16-interactome, we revealed that miR-16 mainly interacts via non-canonical base-pairing to a subset of RNAs, promoting their expression levels (sponge RNAs). Consequently, the canonical miR-16 activity, involved in the RNA decay of oncogenes such as *cyclin D1* and *D3,* is impaired. This miR-16 non-canonical base-pairing to sponge RNAs can explain both the derepression of miR-16 targets and the promotion of oncogenes expression observed for patients with poor overall survival in two cohorts. miR-16 activity assessment using our sponge-signature discriminates the patient’s overall survival as efficiently as the current method based on copy number variations and driver mutations detection. To conclude, miRNA loss of function due to miRNA sequestration seems to promote cancer burden by two combined events – “loss of brake and an acceleration”. Our results highlight the oncogenic role of the non-canonical base-pairing between miRNAs/mRNAs in uveal melanoma.

## Main text

Uveal melanoma (UM) is the most common primary intraocular tumor in adults and no efficient treatment is currently able to counteract UM metastases (1). In 2017, an integrated analysis of 80 primary UMs has been performed by The Cancer Genome Atlas (TCGA) in order to identify the deregulated pathways in this rare cancer and, *in fine*, to uncover druggable targets (2). Four mRNA signatures have been generated based on tumor progression. Other signatures have been published with few common genes (3–5). Unfortunately, no clear link has been made between genes forming these signatures, suggesting that molecular actors driving this cancer are not fully understood.

UM is currently considered as a G-protein-coupled receptor (GPCR) disease with BES (*BAP1*, *EIF1AX* and *SF3B1*) alterations and copy number variations (1,2). Monosomy 3 is clearly associated with a high risk of metastasis. Apart from BAP1 (3p21.1), the contribution of other genes located on chr 3 to the tumor aggressiveness is not elucidated. Because one copy of *MIR16* gene is located on chr 3 (Fig. 1A), we hypothesized that tumor suppressor activity of miR-16 is decreased in UM patients with monosomy 3 (Fig. 1B and S1). Indeed, a genetic alteration in the *MIR16* locus triggers the development of prostatic cancer, pituitary cancer and chronic lymphocytic leukemia (CLL) (6,7).

**Figure 1:**
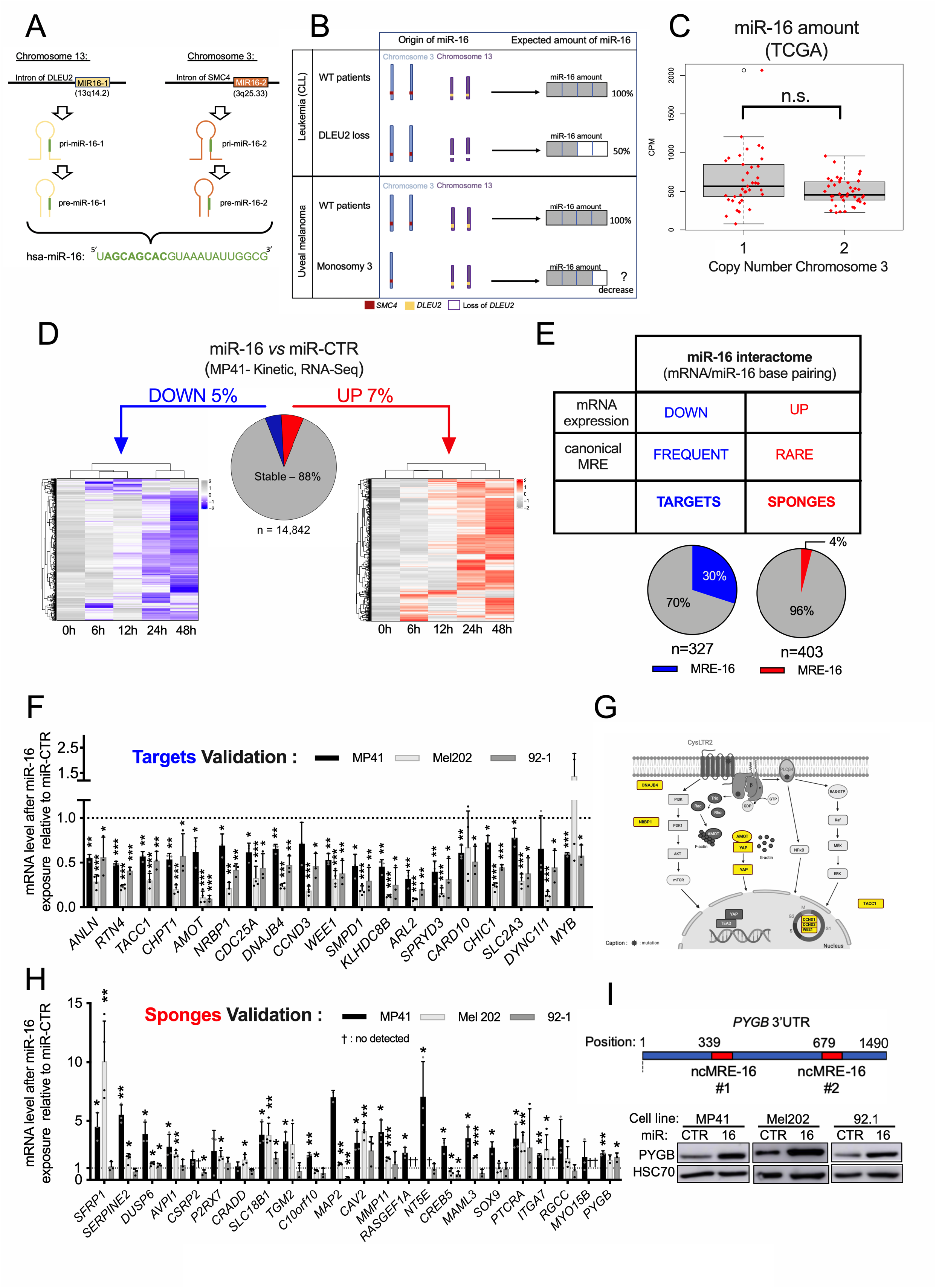
Dual role for miR-16 in uveal melanoma. **A**- Schematic representation of the genomic loci of miR-16 (MIR16-1 & MIR16-2) and miR-16 precursors: pri- & pre-miR-16 (1 & 2) and miR-16. The bolt region on the sequence of miR-16 sequence corresponds to the seed region of the miRNA. **B**- Schematic representation of the expected amount of miR-16 according to the chromosomal status for the two miR-16 loci, for both leukemia and uveal melanoma patients; (CLL: Chronic Lymphocytic Leukemia). **C**- Boxplots of miR-16 expression according to the status of the chromosome 3 for uveal melanoma patients from the TCGA cohort (expressed in counts per million) - monosomy n= 37 and disomy n=42. Each histogram represents the mean ± s.d . n.s. : non significant. **D**- Heatmap representing the differential transcriptomic response induced by transfection of miR-16 *versus* miR-CTR in MP41 cell line (0h= starting time point, 6h, 12h, 24h, 48h post transfection). MP41 transcriptome (n=14,842 genes) are divided in three populations. By comparing miR-16 condition *versus* miR-CTR at early time point (6h-12h) and late time (24h-48h) three populations have been identified: stable genes ~88%, downregulated (LogFC<−0,5) genes ~5% and upregulated (LogFC>0,5) genes ~7%. Left heatmap illustrating the down-regulated RNAs in response to miR-16 transfection in MP41 (0h, 6h, 12h, 24h, 48h). Right heatmap illustrating the up-regulated RNAs (Table S1). **E**- Table describing the expected miR-16 interactome (mRNAs interacting with miR-16) in function of the experimental workflow detailed in Fig. S3A. In function of their expression levels (in response to synthetic miR-16), these miR-16 interacting mRNA have been considered as targets or sponges. MRE for miRNA responsive element. Pie charts indicate the percentage of mRNAs (targets or sponges) 10arbouring at least one MRE-16 have (predicted by TargetScan 7.2) (13). **F**- mRNA expression levels of selected miR-16 targets, 72h after transfection of miR-16 relative to miR-CTR in MP41, Mel202 and 92-1 cells. n=3, 4, 3 biologically independent experiments, respectively. Each histogram represents the mean ± s.d.; Bilateral Student test (with non-equivalent variances) *p<0,05; **p<0;01; ***p<0,001. **G**- Scheme summarized the most frequent genetic alterations found in uveal melanoma and the potential roles of several miR-16 targets in these deregulated pathways (Created with BioRender.com). **H**- mRNA expression levels of selected miR-16 sponges, 72h after transfection of synthetic miR-16 relative to miR-CTR in MP41, Mel202 and 92-1 cells. n=3, 4, 3 biologically independent experiments respectively. Each histogram represents the mean ± s.d.; Bilateral Student test (with non-equivalent variances) *p<0,05; **p<0;01; ***p<0,001; †: Not detected genes. **I**- Schematic representation of the 3’UTR of *PYGB* mRNA containing two non-canonical MREs predicted by RNAHybrid (upper panel). Protein expression levels of PYGB in MP41, Mel202 and 92-1 cell lines in response to miRNA transfection (72h, miR-CTR *versus* miR-16). The picture is representative of n=3, 2, 3 biologically independent experiments, respectively.

We firstly evaluated the expression levels of miR-16-5p (miR-16) relative to chr 3 copy number (Fig. 1C & S2A). Unexpectedly, no significant difference was found between groups of patients in the TCGA cohort. However, we previously showed that miR-16 activity is not always correlated to miRNA expression (8). Sequestration of miR-16 by coding and non-coding RNAs, referred to miRNA-sponges, can dampen the miRNA activity as we demonstrated in cutaneous melanoma. Repression of the miR-16 target mRNAs is thus alleviated, promoting *in fine* tumor growth (9,10).

We hypothesized comparable mechanisms mediate miR-16 inactivation in UM. To test this hypothesis, we first investigated the tumor suppressor activity of miR-16 in UM cells by elevating miR-16 expression levels through transfection of a synthetic miR-16. UM cell density decreased specifically after 72h after transfection of the synthetic miR-16 (Fig. S2B & S2C), suggesting that indeed miR-16 acts as a tumor suppressor in human uveal melanoma. However, miR-16 levels reached after transfection are more important than physio-pathological levels (basal miR-16 is >3000 copies per cell; Fig. S2A) (11), suggesting that the stoichiometry between miR-16 and its target RNAs was not respected. Knowing that a sequestration mechanism would imply a ‘target shift’ characterized by a lower decay of the canonical miR-16 targets (*CCND1*, *CCND3*, *WEE1* mRNAs) (12) and a miR-16 sponging by other RNAs (10), we defined the miR-16 interactome (mRNAs interacting with miR-16 - Fig. S3, 1D-E, and Table S1) by RNA-pull down and we combined the results with a transcriptomic profiling in response to synthetic miR-16 transfection in uveal melanoma cells (MP41). As expected, we identified downregulated RNAs associated with the presence of canonical miR-16 binding sites (predicted MRE-16 in 30% of target RNAs) (13). Interestingly, we identified another set of RNAs, for which the expression levels increased despite miR-16 interaction. The miR-16 base-pairing seemed non-canonical because only 4% of the sponge RNAs exhibited a predicted MRE-16 (Fig. S4 & Fig. 1D-E). We next confirmed the most interesting targets and sponges by RT-qPCR in two additional UM cell lines (Fig. 1F, H). For the majority of tested RNAs, we validated the increase of sponge RNAs in response to miR-16 in at least another cell line. Although a fraction of sponges seemed to be cell-line specific (Fig. 1H), others, like *PYGB*, were upregulated in response to miR-16 in the three models (Fig. 1H). *PYGB* up-regulation was also detected at the protein level (Fig. 1I). In addition, 50% of the tested putative sponge mRNAs were still increased after transfection of miR-16 in DROSHA knock-out cells in which almost all miRNAs including miR-16 are lost (14), suggesting that miR-16 acts directly on these sponges rather than through a competition with another miRNA involved in sponge decay (Fig. S5A, B). A direct effect of miR-16 on sponges was confirmed by luciferase assay using non-canonical sites of *PYGB*, wild-type or mutated, fused with the luciferase coding sequence (Fig. S5C, D).

To further challenge the miR-16 sequestration hypothesis while preserving the stoichiometry between miR-16 and its interactome, we next depleted *PYGB* mRNA and quantified candidate endogenous miR-16 targets (Fig. S5E, F) selected as a function of: (*i*) a miR-16-dependent mRNA decay (Fig. S6A-B), (*ii*) presence of a predicted MRE-16 in their 3’UTR (Fig. 1E & S3C), and (*iii*) a decrease in MP41 cell density in response to their depletion (Fig. S6C, D). Since the depletion of only one miR-16 sponge (*PYGB*) was followed by a moderate decrease of several miR-16 target RNAs (Fig. S5E, F), it is tempting to conclude that miR-16 sequestration involves several RNAs with non-canonical MRE-16. This model of sequestration may explain why we identified 57 potential miR-16 sponges in UM (Fig. S3B). Moreover, our model predicts that miR-16 sequestration by sponges should abolish the canonical activity of miR-16, leading to a derepression of the miR-16 targets involved in cell proliferation and/or survival of UM cells (Fig. 1G & S3B). Thus, we investigated if a high level of miR-16 sponges could be associated with a loss of canonical miR-16 activity and consequently associated with a poor overall survival of patients. We demonstrated that quantification of 57 miR-16 sponge candidates efficiently predicted survival in UM patients (TCGA cohort), reflecting metastasis risk (Fig. 2). Unsupervised gene expression analysis identified 2 clusters: light and dark grey (cluster 2 & 1, respectively) (Fig. 2A). These clusters are highly correlated with those defined by TCGA. Remarkably, miR-16 expression level was comparable in the two groups supporting the sponging hypothesis (Fig 2B). In accordance with our hypothesis, we showed that a high level of miR-16 sponges is associated with a dismal survival (Fig. 2C) and miR-16 targets are derepressed in cluster 1 (Fig. 2C and S7). Altogether, these results indicate that miR-16 activity (appraised using miR-16 sponges and target expression levels) is a useful marker for clinicians in contrast to miR-16 expression. Since 57 RNAs are too many to be exploited clinically, we developed a risk model (5) (Fig. 2D) (Table S2), identifying four RNAs to predict the overall survival of patients with UM (signature S4: Fig. 2D-E & S8). This ability of the S4 signature to predict survival was confirmed in an independent cohort (n=63; GSE22138) (15) (Fig. 2F).

**Figure 2:**
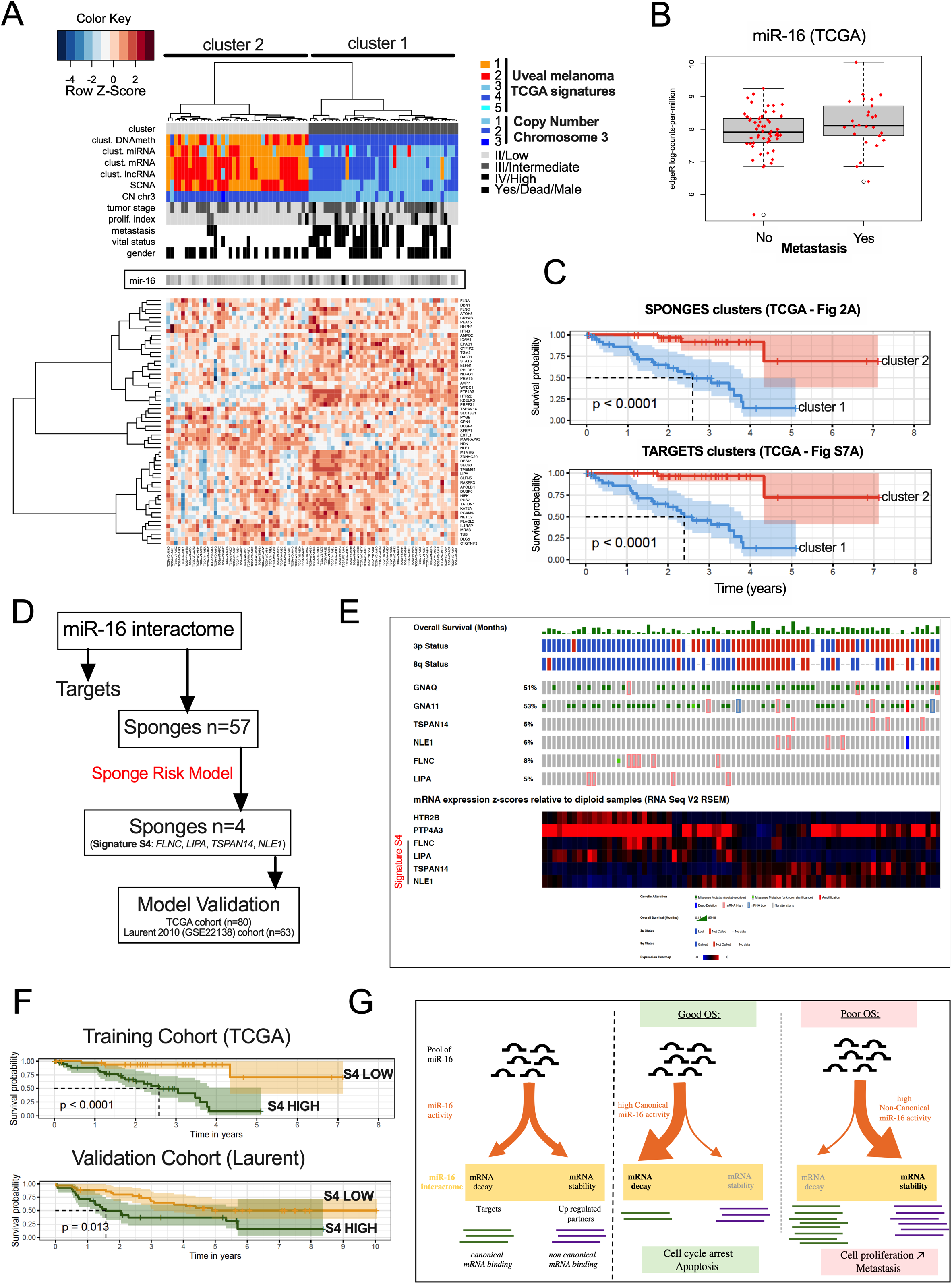
miR-16 availability defines two signatures predicting the prognostic of uveal melanoma patients. **A**- Heatmap depicting the expression levels of 57 sponge RNAs - TCGA cohort of uveal melanoma. Unsupervised gene expression analyses identified 2 clusters: light and dark grey (clusters 2 & 1, respectively). Cluster 1 is associated with poor clinical outcome (chromosome 3 monosomy, metastasis, …). Moreover, cluster 1 overlaps with the TCGA signatures (miRNA, mRNA, lncRNA & DNA methylation) previously associated with poor clinical outcome. **B**- Boxplot representing the amount of miR-16 in function of the metastasis status (TCGA cohort). No significant difference was found. **C**- Determination of overall survival curves by Kaplan–Meier analysis based on clusters 1 & 2 The difference in survival between groups is reported (log-rank test p-value). KM analyses have been performed for miR-16 sponges RNA and targets RNA according to the clusters 1 & 2 defined in Fig. 2A and S7A (Table S1). **D**- The Sponges risk model workflow identifying 4 sponges RNAs (the signature S4). The TCGA-UVM cohort has been used as a training cohort and the GEO dataset GSE22138 as a validation cohort. We trained an optimal multi-gene survival model based on the expression of the sponges in the training cohort by selecting survival-associated genes with the *rbsurv* R package using 1,000 iterations. Briefly, this package allows a sequential selection of genes based on the Cox proportional hazard model and on maximization of log-likelihood (see Methods and Table S2). Risk scores were determined using classical Cox model risk formulae with a linear combination of the gene expression values weighted by the estimated regression coefficients. The risk cutoff was set to the median of the linear predictor. The Kaplan–Meier method was used to estimate the survival distributions. Log-rank tests were used to test the difference between survival groups. Analyses were carried out with the *survival* and *survivalROC* R packages. **E**- Genetic alterations described in the TCGA cohort of uveal melanoma from Cbioportal for our 4 sponges (*TSPAN14*, *NLE1*, *FLNC* & *LIPA*). *GNAQ & GNA11* were used as controls (upper panel). mRNA expression (z-scores, lower panel) of the 4 sponges has been compared to two mRNA highly expressed in patients with a poor clinical outcome (*HTR2B* & *PTP4A3*). The complementarity of the 4 sponges efficiently discriminates the overall survival of the patients (TCGA cohort). **F**- Determination of the overall survival curves by Kaplan–Meier analysis based on the sponge risk model in two cohorts (signature S4). The risk cutoff (low/high) was set to the median of the linear predictor. **G**- Hypothetical molecular mechanism explaining the loss of tumor suppressor activity of miR-16 by RNAs (loss of brake effect). miR-16 is considered as a potent tumor suppressor because it regulates the cell cycle by decreasing the expression level of targets such as *CCND3* and *WEE1*. In patient with a poor OS, miR-16 is not able to bind and regulate these RNAs. The sequestration of miR-16 on other mRNAs (defined as sponges) is associated with metastasis risk and dismal overall survival in UM. Instead of promoting RNA decay of miR-16 targets, the non-canonical miR-16 activity promotes expression of sponges such as the pro-tumoral PTP4A3 gene (acceleration effect). miR-16 sequestration seems to promote cancer burden by two combined events – “loss of brake and an acceleration”. In conclusion, we propose that miR-16 can exert pro- or anti-tumoral activity in function of its base-pairing to mRNAs. For clinicians, our signature S4 accurately predicts clinical outcomes compared with existing classification schemes. Our results expand the current knowledges on molecular mechanisms promoting uveal melanoma and pave the way to explore new therapeutic candidates targeting miR-16 activity for a cancer without effective treatment at metastatic stage.

## Conclusion

Here, we characterized a molecular mechanism explaining the loss of tumor suppressor activity of miR-16 by RNAs (loss of brake effect), which is associated with metastasis risk and dismal overall survival in UM (Fig. 2G). Instead of promoting RNA decay of miR-16 targets, the non-canonical miR-16 activity mediates the expression of pro-tumoral genes such as *PTP4A3* (acceleration effect) (15).

## Discussion

Although the concept of competition between RNAs remains a matter of debate (16) the factors regulating the balance between miR-16 canonical and non-canonical activity (17) may represent new vulnerabilities that could be targeted to treat UM, a GPCR & miR-16 disease.

## ABBREVIATIONS

UM: Uveal melanoma
TCGA: the cancer genome atlas
GPCR: G-protein-coupled receptor
BES: *BAP1*, *EIF1AX* and *SF3B1*
BAP1: BRCA1 associated protein-1
EIF1AX: Eukaryotic Translation Initiation Factor 1A X-Linked
SF3B1: Splicing Factor 3b subunit 1
CLL: chronic lymphocytic leukemia (CLL)
MRE: MicroRNA Recognition Element
CCND1: Cyclin D1
CCND3: Cyclin D3
PYGB: Glycogen phosphorylase B
DROSHA: Drosha ribonuclease III
PTP4A3: Protein tyrosine phosphatase type IVA, member 3.

## ACKNOWLEDGEMENTS

The authors thank the Gene Expression and Oncogenesis team from the CNRS UMR6290, Dr Pascal Loyer from NuMeCan (INSERM U1241), BIOSIT core facilities of Rennes 1 University (SFR UMS CNRS 3480 – INSERM 018, especially P. Gripon for the BSL3), the UCA GenomiX platform of IPMC and the Centre de Ressources Biologiques humaines Santé (especially C. Pangault) for their help. The authors thank Dr FA Karreth, Dr M. Migault, Pr MH Stern, Sylvain Martineau and Pr Stéphan Vagner for helpful discussion. Support was provided by a “Ligue Nationale Contre le Cancer” (LNCC) fellowship and French Ministry of Research (“Ministère français de lʼEnseignement supérieur, de la Recherche et de lʼInnovation”) fellowship (AQ). The authors are grateful to Narry Kim for providing the HCT116 KO DROSHA and HCT116 WT (Korean Collection for Type Cultures (KCTC)) and to Didier Decaudin for the MP41 cell line.

## AUTHOR CONTRIBUTIONS

Conceptualization: AQ & DG.

Methodology: AQ, LB, MA, SA & DG.

Software: AQ, MA, SA, GS & DG.

Formal analysis: AQ, LB, MA, SA, DL & BM.

Investigation: AQ, LB, MA, FC, DL & DG.

Ressources : MA, SA, GS, DD & BM.

Writing-original draft: AQ, LB, FM & DG.

Writing – review & editing : all authors.

Visualization: AQ, LB, MA, SA, GS, BM & DG.

Supervision: DG.

Project Administration: DG & MDG.

Funding: DG & MDG.

## AVAILABILITY of DATA and MATERIALS

Further information and requests for resources and reagents should be directed to and will be fulfilled by David Gilot. All unique/stable reagents generated in this study are available from the Lead Contact with a completed Materials Transfer Agreement. mRNAseq and RIPseq data that support the findings of this study have been deposited in the Gene Expression Omnibus (GEO) under accession code GSE180399 (https://www.ncbi.nlm.nih.gov/geo/query/acc.cgi?acc=GSE180399) and ArrayExpress under accession code E-MTAB-10940 (*https://www.ebi.ac.uk/arrayexpress/experiments/E-MTAB-10940*).

## ETHICAL APPROVAL AND CONSENT TO PARTICIPATE

Not applicable.

## CONSENT for PUBLICATION

Not applicable.

---

**Figure S1:**
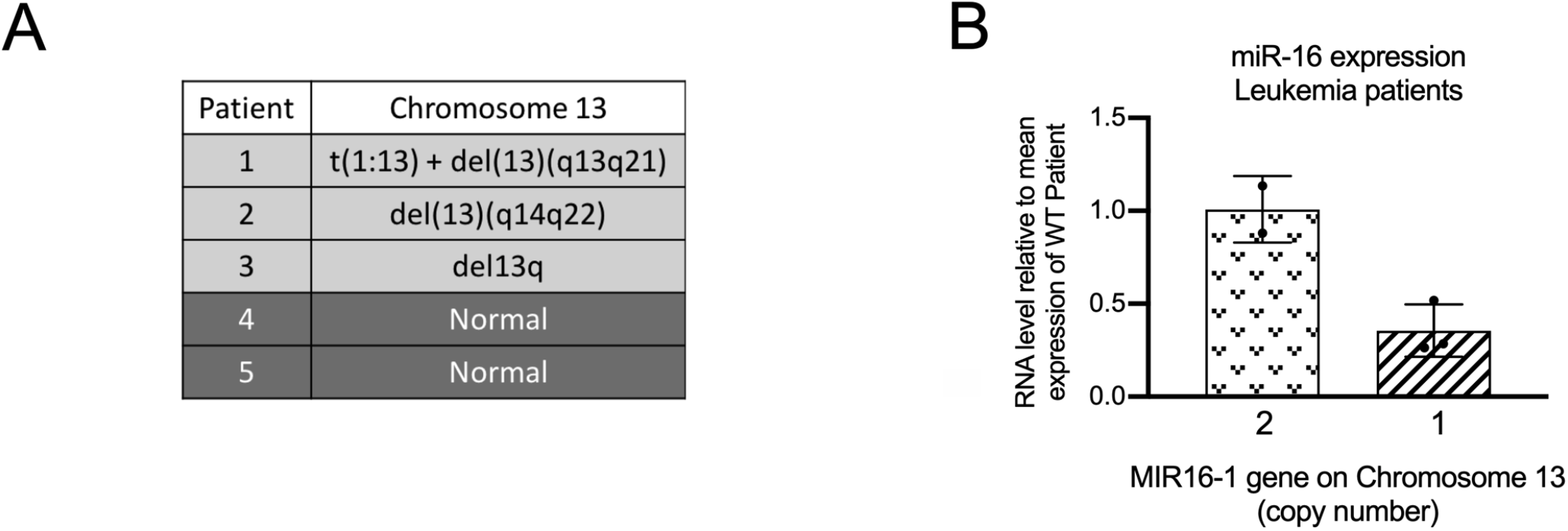
loss of MIR16-1 gene decreases miR-16 expression in patients with leukemia. **A**- Table of cytogenetic features of the chromosome 13 in patients analysed in S1B; del: deletion (n=3); normal (n=2). **B**- miR-16 expression in 5 leukemia patients with or without deletions on chromosome 13. Each histogram represents the mean ± s.d.

**Figure S2:**
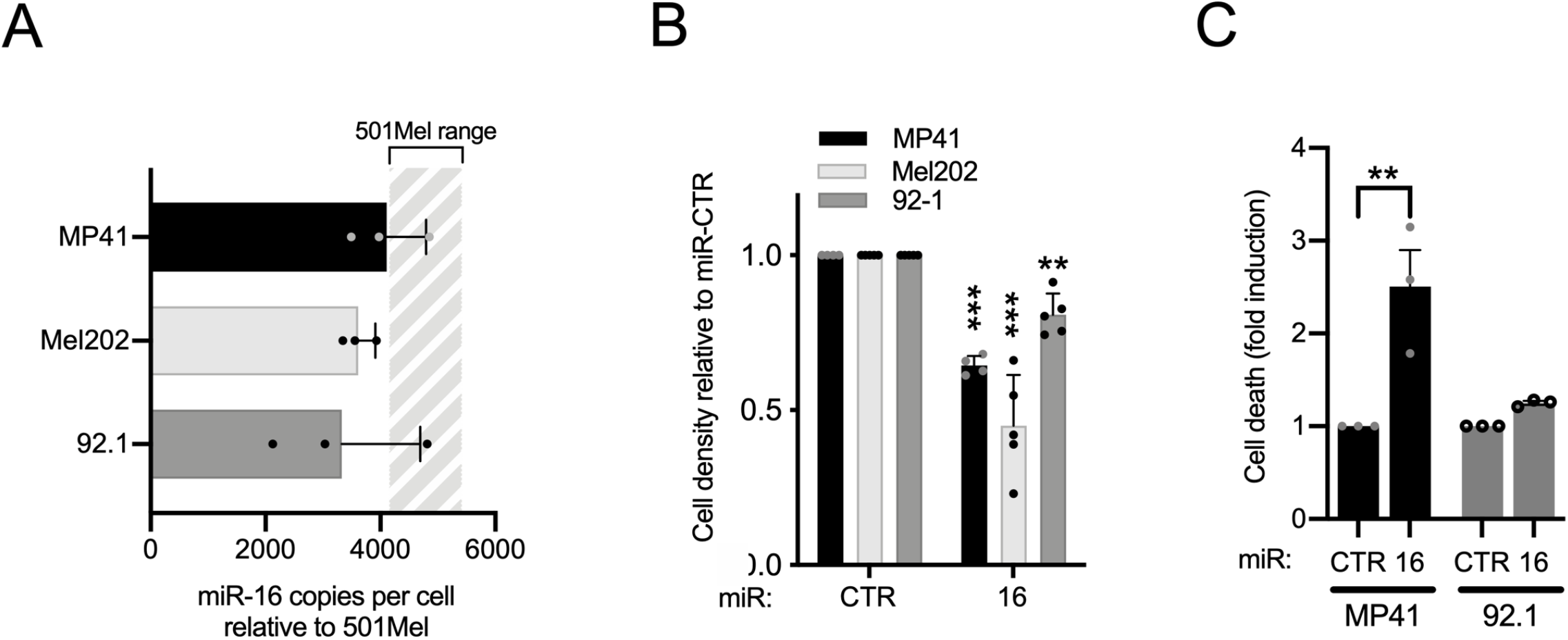
miR-16 expression levels and effects in uveal melanoma cell lines. **A**- Quantification of miR-16 expression in three uveal melanoma cell line (MP41, Mel202, 92-1) by RT-qPCR. Levels were compared to values previously quantified in cutaneous melanoma cell line (501Mel). The absolute quantification (copy number) of miR-16 in 501Mel was determined by Northern-blot (1). n= 3 biologically independent experiments for each cell line. Each histogram represents the mean ± s.d. **B**- Cell density of MP41, Mel202 and 92-1 in response to miR-16 overexpression (transfection of synthetic miR-16 *versus* miR-CTR), 72h after transfection. n=4, 5 and 5 biologically independent experiments, respectively. Each histogram represents the mean ± s.d.; Bilateral Student test (with non-equivalent variances) **p<0;01; ***p<0,001. **C**- Fold induction of dead cells (apoptosis + necrosis; in % relative to miRNA control) in response to the miR-16 overexpression in MP41 and 92.1 cells, 72h after transfection. n=3 biologically independent experiments. Each histogram represents the mean ± s.d.; Bilateral Student test (with non-equivalent variances) **p<0,01.

**Figure S3:**
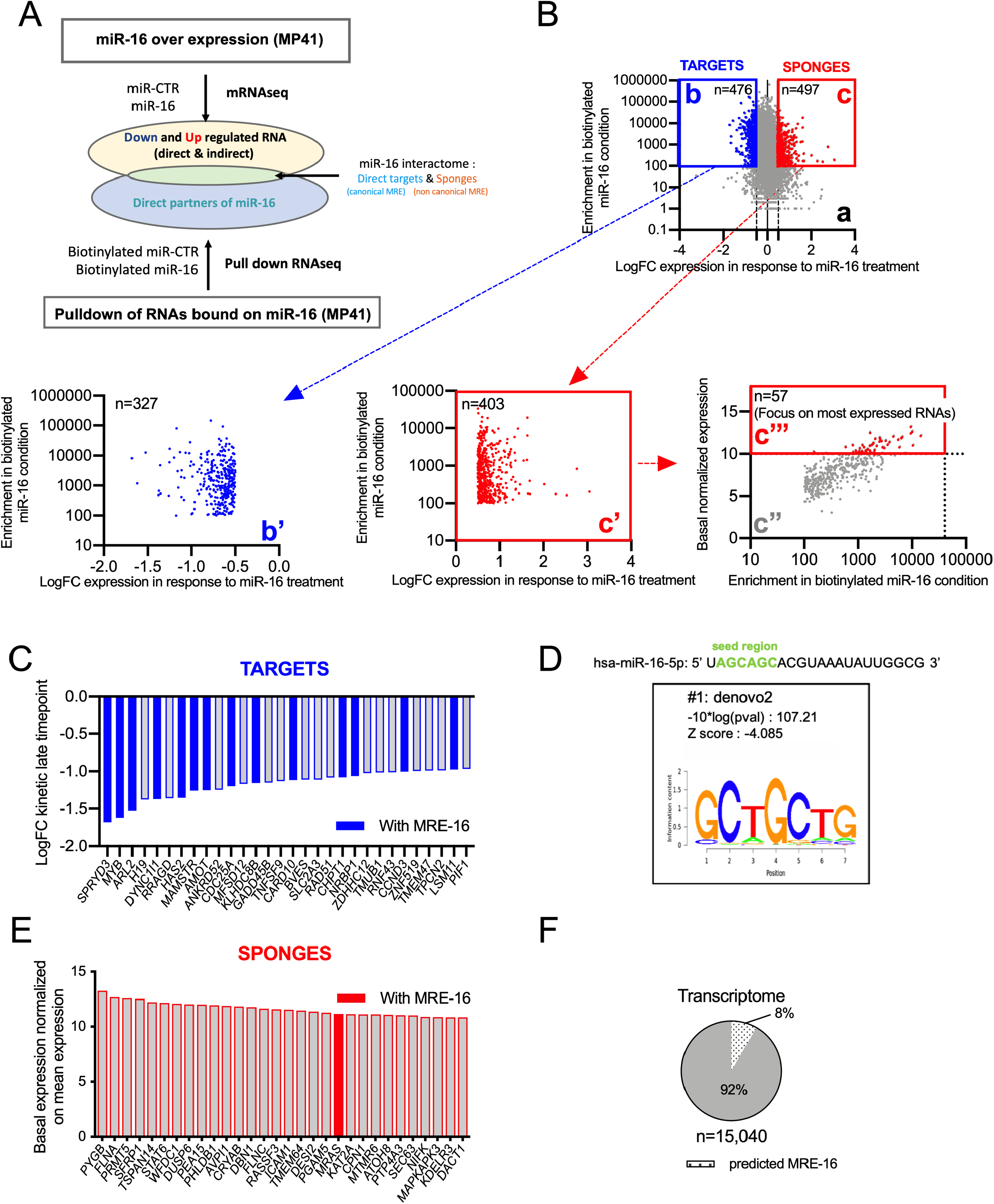
Workflow to uncover miR-16 interactome (targets and sponges) **A**- Schematic representation of the workflow. The kinetic experiment identified 2 RNA populations: downregulated and upregulated RNAs. miR-16 interacting mRNAs have been purified and sequenced using biotinylated miR-16 *versus* biotinylated miR-CTR. This combination of methods identified down- and up- regulated RNAs (targets & sponges) which bind to miR-16. These RNAs defines the miR-16 interactome. B- <u>Graph a</u>: Representation of pulldown enrichment (miR-16 – miR-CTR) in function of the transcriptomic expression changes (Fold change (FC) expression after miR-16 exposure). Two clusters are delimited. The cluster b, in blue, corresponds to RNAs with an enrichment >100 and down regulated with a fold change < −0,5 (n=476 genes). The cluster c, in red, corresponds to RNAs with an enrichment >100 and up regulated with a fold change >0,5 (n=497 genes). <u>Graph b’</u>: represents same genes of the graph b without those suspected to be false positive due to their detection with biotinylated miR-CTR (threshold 500 reads in the RNAseq, for miR-CTR) (n=327 genes). <u>Graph c’</u>: represents same genes of the graph c without those suspected to be false positive due to their detection with biotinylated miR-CTR (threshold 500 reads in the RNAseq for miR-CTR) (n=403 genes). <u>Graph c’’ in grey:</u> is the same selected genes of the graph c’ but they are represented according to their basal expression by pulldown enrichment (miR-16 – miR-CTR). The c’’’ cluster represents only genes with a basal expression >10 (normalized expression). This workflow identified 57 potential sponges. **C**- Downregulated RNAs (miR-16 targets) are ordered according to the level of the downregulation at the late timepoint. Only the top 30 targets are represented. Blue ones harbor at least one canonical MRE-16 predicted by TargetScan 7.2. **D**- Logo of miRNA binding site motifs enriched in cluster down regulated RNAs after miRNA pulldown in MP41 (analyzed by Cistrome SeqPos (2)) **E**- Basal expression of the 30 most expressed genes at the basal level (from the selected genes represents in the graph c’’’ (Fig. S3B)). Red ones harbor at least one canonical MRE-16 predicted by TargetScan 7.2. **F**- Pie chart represents the percentage of RNAs harboring at least one MRE-16 predicted by TargetScan7.2 in the entire MP41 transcriptome, in white (n=15,040)

**Figure S4:**
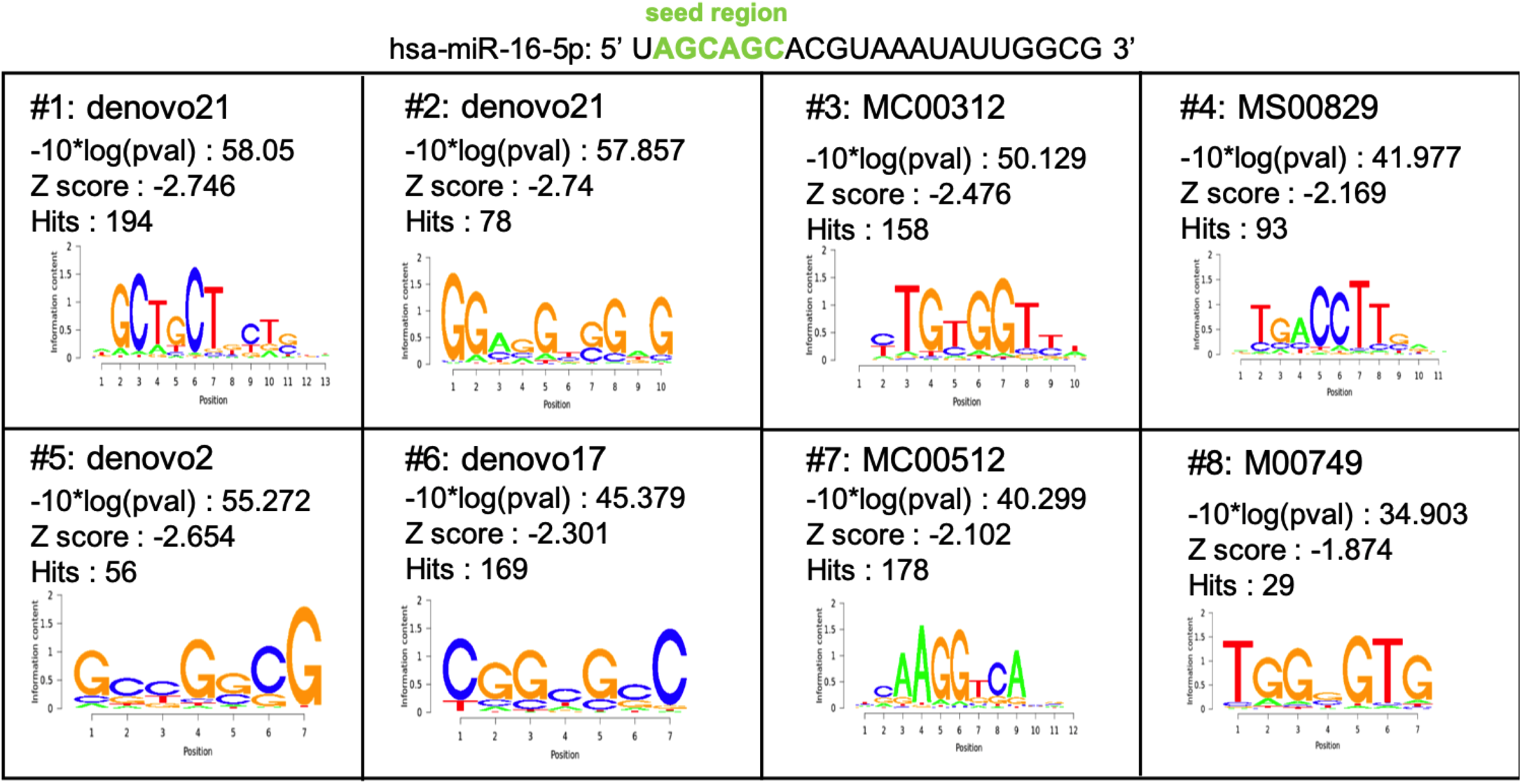
Putative non-canonical miR-16 binding sites on miR-16 sponges. Logos of miRNA binding site motifs enriched in cluster of upregulated miR-16 interactants after miR-16 pulldown in MP41. Analyses were performed using Cistrome SeqPos.

**Figure S5:**
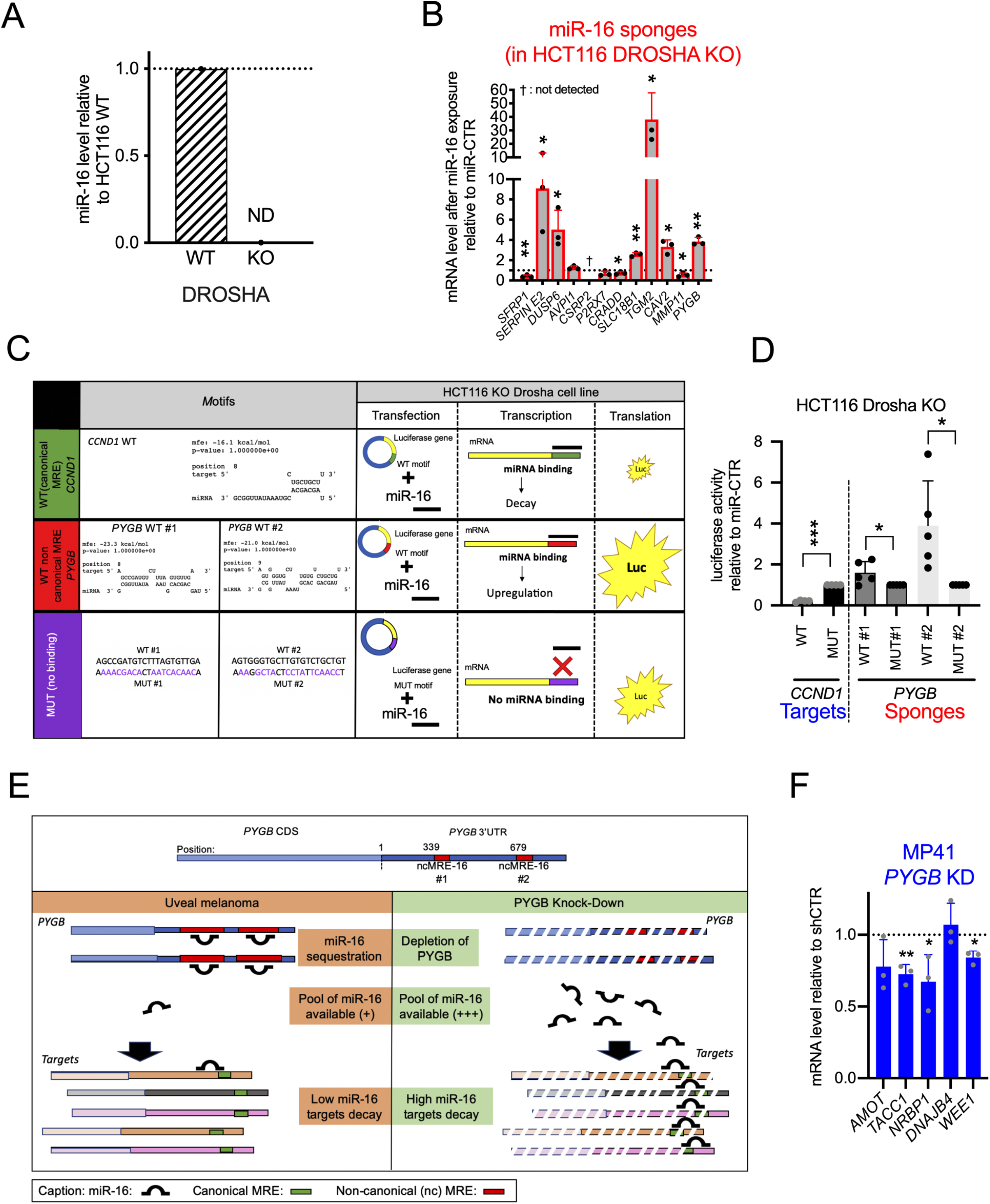
miR-16 is sequestered on non-canonical miR-16 binding sites. **A**- Relative expression levels of miR-16 in HCT116 WT and DROSHA KO assessed by RT qPCR (n=2 biologically independent experiments). **B**- mRNA expression levels of selected miR-16 sponges 72h after transfection of synthetic miR-16 relative to miR-CTR in HCT116 DROSHA knock-out cells. n=3 biologically independent experiments Each histogram represents the mean ± s.d.; Bilateral Student test (with non-equivalent variances) *p<0,05; **p<0;01; †: Not-detected genes. **C**- Biological function of non-canonical MRE-16. On the left side: predicted base-pairing between *PYGB* mRNA and miR-16 using RNAhybrid (3). Base-pairing has been evaluated for wild-type (WT) MRE-16 (non-canonical MRE-16 #1 & #2) from *PYGB* mRNA. On the right side: schematic representation of the luciferase assay. Canonical MRE-16 (from CCND1, (4)) has been used as positive control (miR-16 induced the decay of a mRNA harbouring a canonical MRE-16). The two non-canonical miR-16 binding sites of *PYGB* have been cloned in fusion with the luciferase coding sequence. The translation efficiency of these chimeric RNAs is estimated by assessing the luciferase activity. **D**- Luciferase assay assessing the effect of synthetic miR-16 on these chimeric RNAs in HCT116 KO DROSHA cell line. Canonical MRE-16 (from CCND1, (4)) has been used as positive control (as attended miR-16 induced the decay of a mRNA harbouring a canonical MRE-16). MUT: mutated; WT: wild type (n= 5 biologically independent experiments). Each histogram represents the mean ± s.d.; Bilateral Student test (with non-equivalent variances) *p<0,05. **E**- Hypothetical scheme explaining the miR-16 displacement is response to miR-16 sponge depletion. Sequestered miR-16 on *PYGB* mRNA are released from *PYGB* and reached other miR-16 binding sites (on other RNAs including miR-16 targets). Based on our hypothesis, the expression levels of these targets should thus decrease. *PYGB* mRNA has been selected because it is the most expressed sponge identified in this study. Here, the stoichiometry between miR-16 and miR-16-interacting RNAs is preserved (no miR-16 transfection). **F**- mRNA expression levels of miR-16 targets: *AMOT, TACC1, NRBP1, DNAJB4* and *WEE1* in response to *PYGB* mRNA depletion in MP41 (shPYGB relative to shCTR, (n=3 biologically independent experiments each) Each histogram represents the mean ± s.d.; Bilateral Student test (with non-equivalent variances) *p<0,05; **p<0;01.

**Figure S6:**
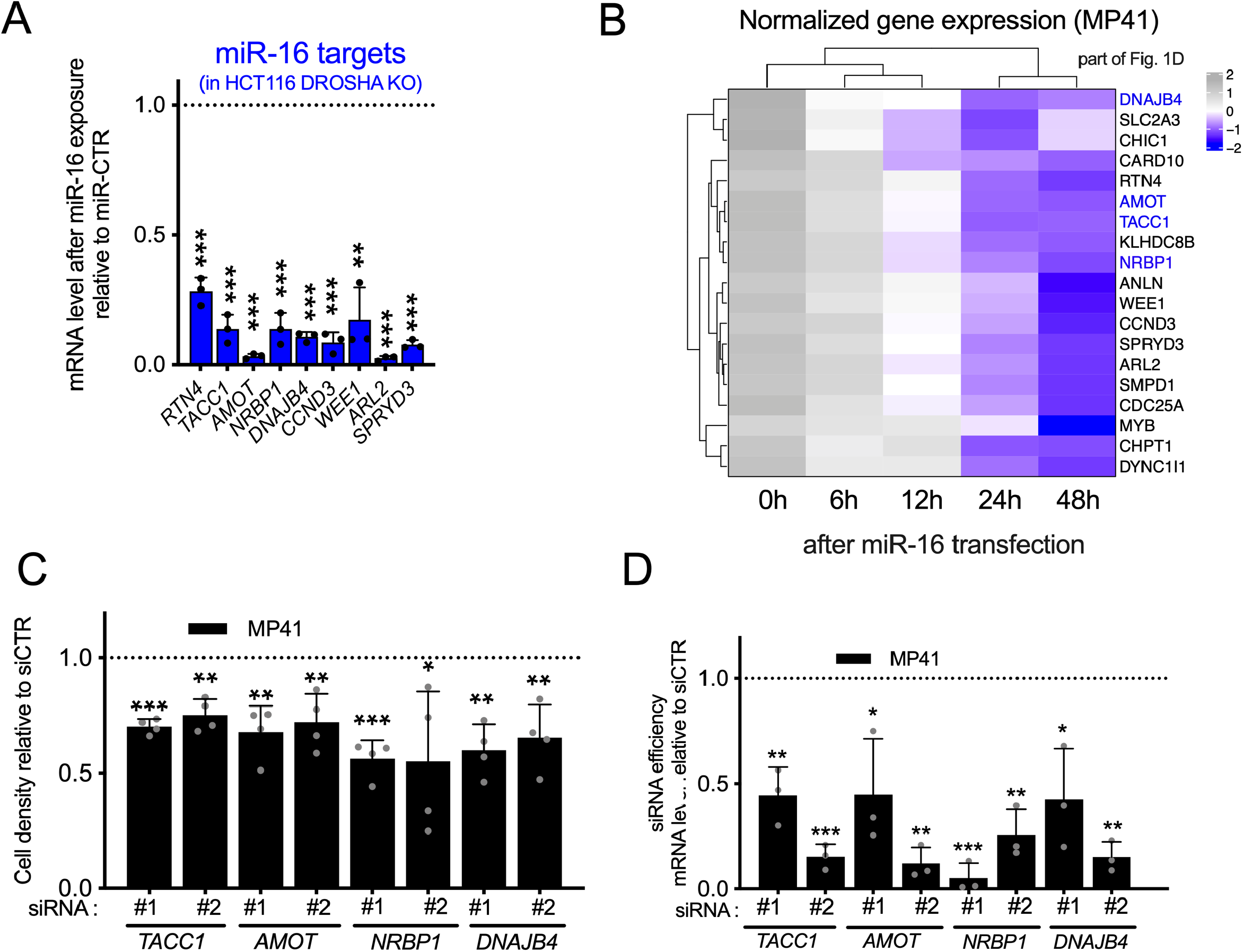
miR-16 modulates cell fate by targeting several RNAs in uveal melanoma. **A**- mRNA expression levels of selected miR-16 targets, 72h after transfection of miR-16 or miR-CTR in HCT116 DROSHA KO. n=3 biologically independent experiments. Each histogram represents the mean ± s.d.; Bilateral Student test (with non-equivalent variances) **p<0,01; ***p<0,001. **B**- part of the Fig. 1D; heatmap illustrating the selected genes of the Fig. 1F, analysed by RNAseq, after miR-16 transfection in MP41 cells (kinetic : 0h, 6h, 12h, 24h, 48h). **C**- Cell density of MP41 cells in response to the depletion of *TACC1*, *AMOT*, *NRBP1*, *DNAJB4* by two different siRNAs (#1 and #2) relative to siCTR, (n=4, biologically independent experiments) 72h after transfection. Each histogram represents the mean ± s.d.; Bilateral Student test (with non-equivalent variances) *p<0,05; **p<0;01; ***p<0,001. **D**-Efficiency of siRNA used for Fig. 6C (evaluated by RT-qPCR, 72h post transfection). Two different siRNA (#1 and #2)/gene. n=3, biologically independent experiments. Expression relative to siCTR, quantified by RT-qPCR 72h after transfection, *P<0,05; **P<0;01; ***P<0,001

**Figure S7:**
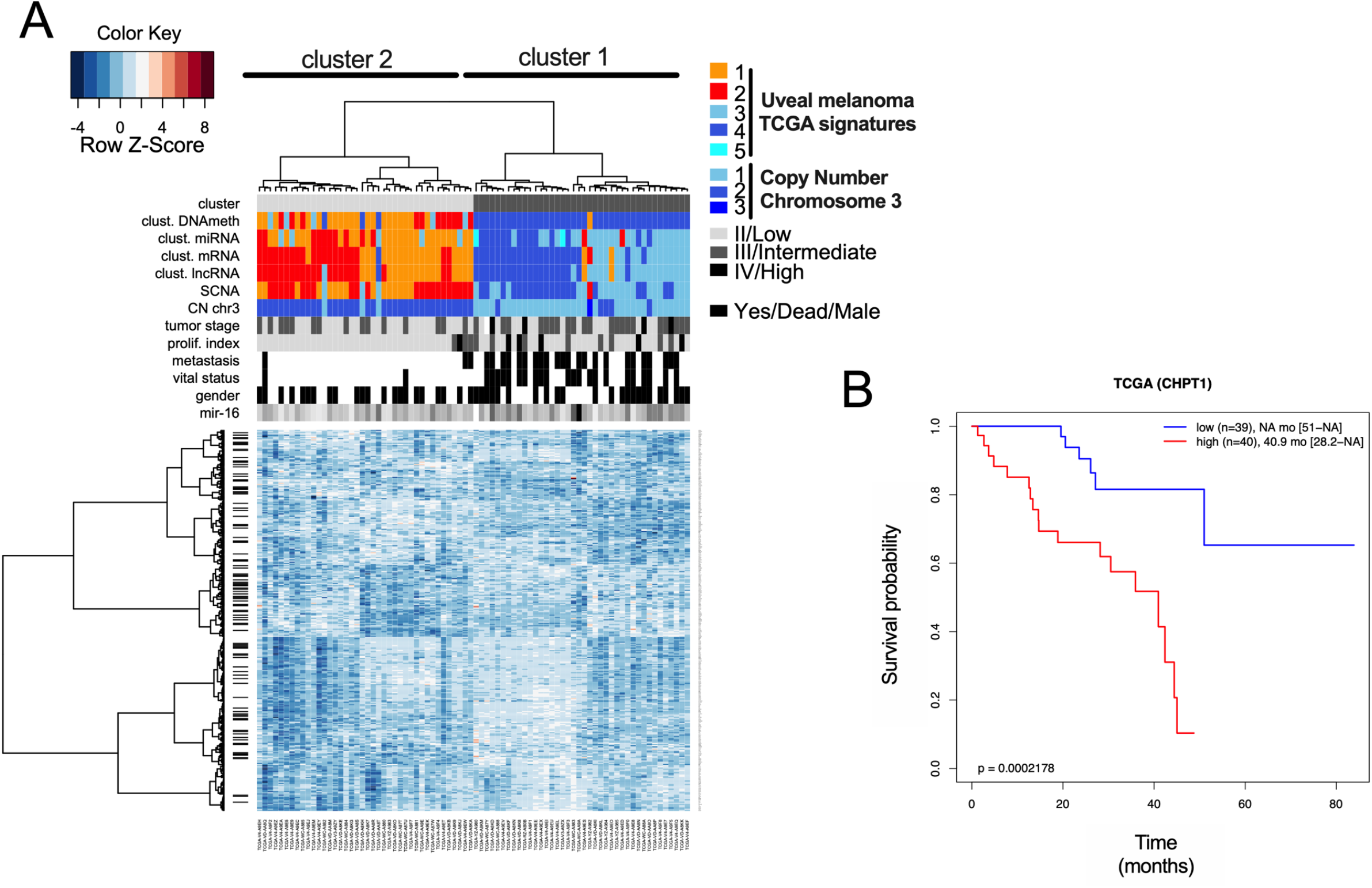
Survival analysis based on miR-16 targets expression. **A**- Heatmap depicting the expression levels of 327 miR-16 targets - TCGA cohort of uveal melanoma. Unsupervised gene expression analyses identified 2 clusters: light and dark grey (clusters 2 & 1, respectively). Cluster 1 is associated with poor clinical outcome (chromosome 3 monosomy, metastasis, …). Moreover, cluster 1 overlaps with the TCGA signatures (miRNA, mRNA, lncRNA & DNA methylation) also previously associated with poor clinical outcome. **B**- Determination of overall survival curves by Kaplan–Meier analysis based on *CHPT1* expression (a miR-16 target) in the TCGA cohort (below or above median expression of the gene). The difference in survival between groups is reported (log-rank test p-value). *CHPT1* has been selected to illustrate the fact that a high expression level of sponges is associated with a high level of miR-16 targets as illustrated in Fig. 2G.

**Figure S8:**
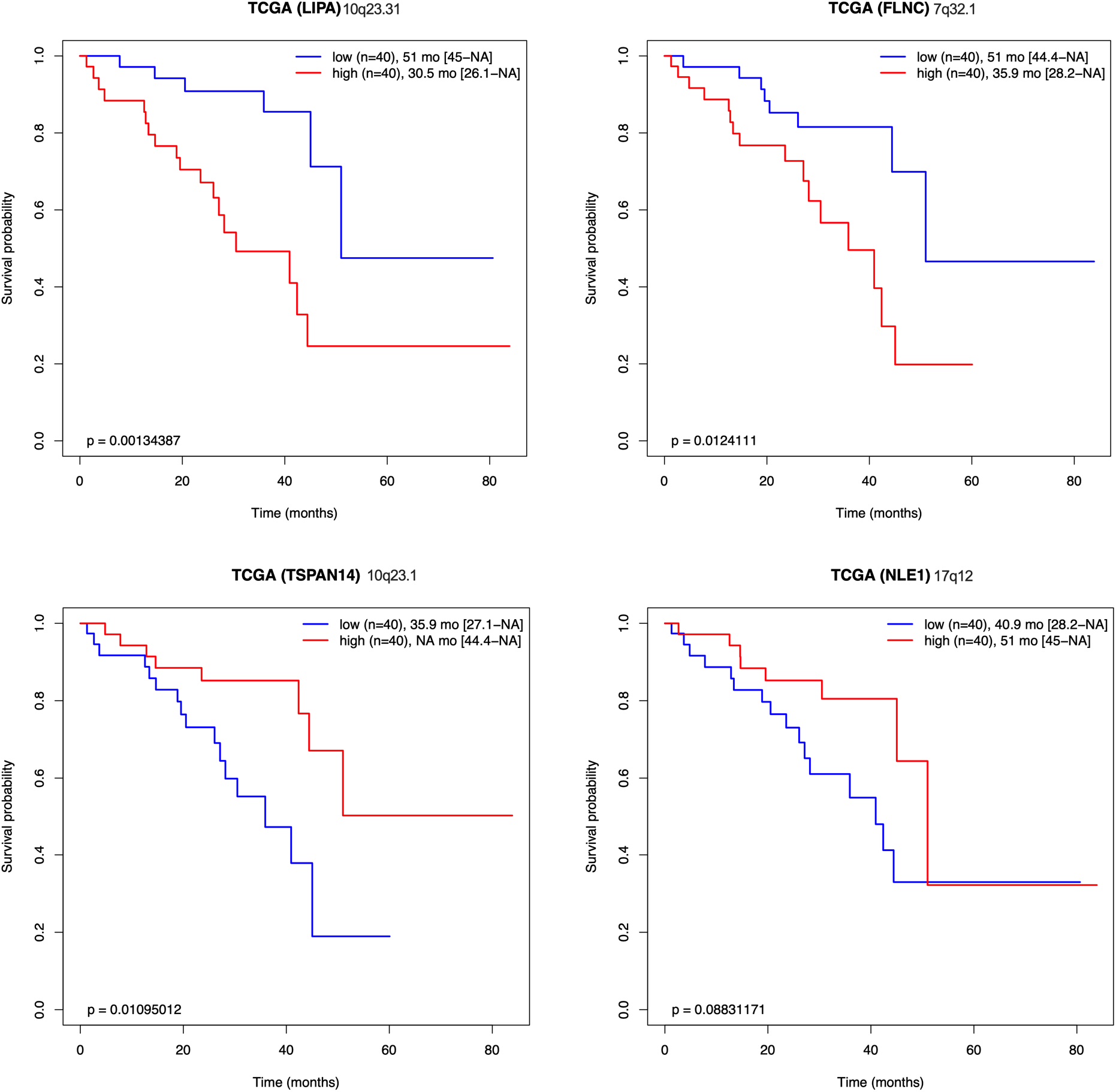
Survival analysis based on signature S4 genes. **A**- Kaplan–Meier analyses for each sponge from the Signature S4 (*LIPA*, *FLNC*, *TSPAN14* & *NLE1*). Overall survival after subdivision into low (blue) and high (red) expression groups (below or above median expression of the gene). The size and the median survival of each group are specified (with 95% CI between brackets). The difference in survival between groups is also reported (log-rank test p-value). Chromosome position is specified for each sponge.

