## Supplementary data for "Uveal Melanoma: a miR-16 disease?"

### SUPPLEMENTARY INFORMATION & EXPERIMENTAL PROCEDURES

Four tables and eight figures.

**Table S1: miR-16 interactome and signatures**

**Table S2: Sponges Risk Model**

**Table S3: Sequences of primers, siRNAs and antibodies**

**Table S4: Raw data**

### EXPERIMENTAL PROCEDURES

Cell Lines and Culture Conditions

MP41 cell line was obtained from Decaudin's lab at Curie Institute, Paris, France. Mel202 and 92.1 cell line were obtained from European Collection of Authenticated Cell Cultures (ECACC) (Merck). 501Mel cell line were obtained from American Type Culture Collection (ATCC). HCT WT and KO DROSHA cell lines were obtained from Korean Collection for Type Cultures (KCTC), Microbial Resource Center. All cell lines were maintained in humidified air (37°C, 5% CO2). Uveal melanoma cell lines were maintained in RPMI-1640 medium (Gibco™, ThermoFisher, MA) supplemented with 20% Fetal Bovine Serum (FBS) (EurobioScientific) and 1% Penicillin-Streptomycin (PS) antibiotics (Gibco™, ThermoFisher). 501Mel were maintained in RPMI-1640 medium (Gibco™, ThermoFisher) supplemented with 10% Fetal Bovine Serum (FBS) (EurobioScientific) and 1% Penicillin-Streptomycin (PS) antibiotics (Gibco™, ThermoFisher). HCT116 cell lines were maintained in McCoy's 5A (Gibco™, ThermoFisher) supplemented with 10% Fetal Bovine Serum (FBS) (EurobioScientific) and 1% Penicillin-Streptomycin (PS) antibiotics (Gibco™, ThermoFisher). All cell lines have been routinely tested for mycoplasma contamination (Mycoplasma contamination detection kit; rep-pt1; InvivoGen).

siRNAs and miRNA transfection

Sequences are available in Table S3. All siRNAs and synthetic mimics were transfected at 66nM using Lipofectamine RNAiMAX (ThermoFisher Scientific) according to manufacturer's instructions. For cell density assay: 80,000 cells for 92-1 and HCT116 KO Drosha, HCT116 WT and 10,000 for MP41 and ; 12,000 for 92-1 cells were seeded in 96-well plates, in quadruplicates. For RNA and proteins analysis, 250,000 cells were seeded in 6-well plates. Cells were harvested 72h after transfection (or kinetic). All siRNAs were purchased from IDT DNA. All mimics were purchased from Dharmacon.

shRNA experiments

Lentiviral particles carrying shRNA vectors targeting human *PYGB* mRNA (shPYGB, TL310025V), and scramble shRNA (shCTR, TR30021V) were purchased from Origen. Lentiviral production was performed as recommended (http://tronolab.epfl.ch) using HEK

293T cells. After infection, cells were maintained in the presence of puromycin 1 µg/ml for selection (Invivogen).

<u>RNA and miRNA isolation, Reverse Transcription, quantitative PCR</u>

RNA was isolated from cell samples using NucleoSpin RNA Plus kit (Macherey-Nagel) and quantified using a NanoDrop 1000 Spectrophotometer (ThermoFisher Scientific). Reverse transcription was performed with the High Capacity cDNA Reverse Transcription kit (Applied Biosystems). Quantitative PCR was performed on 1 ng cDNA, in 384-well plates using the SYBR$^{\text{TM}}$ Green PCR Master Mix (Applied Biosystems) with the QuantStudio$^{\text{TM}}$ 7 Flex Real-Time PCR System (Applied Biosystems). RNA levels were normalized against human *GAPDH*. Relative amounts of transcripts were determined using the $\Delta\Delta - \mathrm{Ct}$ method and human *GAPDH* transcript level was used as an internal control for each cell line sample.

MicroRNA was isolated using mirVana$^{\text{TM}}$ miRNA isolation kit (Ambion, Life Technologies,). Reverse transcription was performed with the TaqMan microRNA Reverse Transcription kit (Applied Biosystems) with the Megaplex RT Primers Pool A v2.1 (Applied Biosystems). Quantitative PCR was performed on 2.5 ng cDNA, in 384-well plates using the TaqMan$^{\text{TM}}$ Gene Expression Master Mix (Applied Biosystems, Foster City, CA) with the QuantStudio$^{\text{TM}}$ 7 Flex Real-Time PCR System (Applied Biosystems, Foster City, CA). Relative amounts of transcripts were determined using the $\Delta\ \Delta - \mathrm{Ct}$ method and human RNU6B was used as an internal control for each cell line sample. The primers were used are described in Table S3.

<u>Cell-density evaluation</u>

Cell density was measured by methylene blue colorimetric assay (Gilot et al., 2017.). Briefly, cells were fixed for 30 min with 70% ethanol. Then, fixed cells were dried and stained 30 min with 1% methylene blue dye in borate buffer. Plates were washed 3 times with fresh tap water and 100 µl of 0.1N HCl per well were added. Plates were analyzed with a spectrophotometer at 620nm.

#### Western-blot experiments

Experiments were performed as previously described (1). Membranes (GE HealthCare, Chicago, IL) were probed with suitable antibodies and signals were detected using the LAS-3000 Imager (Fuji Photo Film). The antibodies are described in Table S3.

#### RNA sequencing

Total RNAs were quantified using a NanoDrop 1000 Spectrophotometer (Thermo Scientific) and RNA integrity (RIN > 8) was evaluated using RNA nano-chips on the Agilent 2100 Bioanalyzer Instrument (Agilent Technologies). Libraries generation and sequencing experiments have been conducted as previously reported (5).

#### Biotinylated miRNA pull down

These experiments were performed on MP41 cell according to the protocol published by Judy Lieberman's lab (6) but with minor modifications. 15 millions of MP41 cells were seeded in 3 X 150mm dishes 5 millions each) and were transfected the next day with 100nM of biotinylated miR-16 or miR-CTR (Dharmacon). Next, they were harvested ~24h post transfection. Cells from the 3 dishes were treated separately. Meanwhile, magnetic beads (Streptavidine Dynabeads™ M-280 DYNAL™, ThermoFisher Scientific) were washed and blocked according to the protocol (6). Cell lysate and washed beads were incubated for 4 hours at 4°C on a rotating agitator. All next steps: the precipitation and the purification of coupled RNA was performed according to the protocol. We pooled the 3 same conditions (from the 3 transfected plates) in the end of the RNA purification. Purified RNA was quantified using quantified using a NanoDrop 1000 Spectrophotometer (ThermoFisher Scientific) followed by the sequencing.

#### Biotinylated miRNA pull down sequencing

RNA sequencing of pull downed RNA has been done by Novogene according to its RIP sequencing protocol (Illumina PE150/Q30≥80%).

#### In silico analyses

The miRNA binding sites on RNA (MRE) were predicted by webtool TargetScan 7.2 (7) and RNAhybrid (3) both available online. Non-canonical MREs have been identified using RNAhybrid and Cistrome SeqPos motif analysis (2). From RNAseq data, 903 genes were found to be down-regulated (with fold change 1.5). miR-16 peaks falling within those genes were called following the procedure described by Sérandour et al. (2012) (8). The resulting bed file containing 504 peaks was used for de novo motif search with the SeqPos tool from Cistrome (2), which looked for enriched DNA motifs within these DNA regions. Annotated genes associated with de novo motifs were identified.

To assess the survival prognosis capabilities of the (selected genes or sponges/targets), we performed univariate Cox analyses of the expression data for these genes, with overall survival (OS) as a dependent variable. Patients were divided into two categories according to the median expression of each gene: low expression (below median) and high expression (above median). The Kaplan–Meier method was used to estimate the survival distributions. Log-rank tests were used to test the difference between survival groups. Analyses were carried out with the survival R package.

Sponge Risk model

We used the TCGA-UVM cohort downloaded from the Xena Browser as a training cohort and the GEO dataset GSE22138 as a validation cohort. We trained an optimal multi-gene survival model based on the expression of the sponges in the training cohort by selecting survival-associated genes with the rbsurv R package using 1,000 iterations. Briefly, this package allows a sequential selection of genes based on the Cox proportional hazard model and on maximization of log-likelihood. To increase robustness, this package also selects survival-associated genes by repetition (1,000 times) of a separation between the training and validation sets of samples. Risk scores were determined using classical Cox model risk formulae with a linear combination of the gene expression values weighted by the estimated regression coefficients. The risk cutoff was set to the median of the linear predictor. The Kaplan–Meier method was used to estimate the survival distributions. Log-rank tests were used to test the difference between survival groups. Analyses were carried out with the survival and survivalROC R packages.

Statistics and reproducibility

Data are presented as mean ± s.d. unless otherwise specified, and differences were considered significant at a P value of less than 0.05. Comparisons were performed using Bilateral Student test (with non-equivalent variances). All statistical analyses were performed using Prism 8 software (GraphPad, La Jolla, CA, USA) or Microsoft Excel software

Overall survival was estimated using the Kaplan–Meier method. Univariate analysis using the Cox regression model or log-rank test, as specified, was performed to estimate hazard ratios (HRs) and 95% confidence intervals (CIs). All experiments were performed three or more times independently under similar conditions, unless otherwise specified in the figure legends (raw data available in Table S4).

Data availability

mRNAseq and RIPseq data that support the findings of this study have been deposited in the Gene Expression Omnibus (GEO) under accession code GSE180399 (https://www.ncbi.nlm.nih.gov/geo/query/acc.cgi?acc=GSE180399) and ArrayExpress under accession code E-MTAB-10940 (*https://www.ebi.ac.uk/arrayexpress/experiments/E-MTAB-10940*), respectively.

The human melanoma data set (uveal melanoma, IlluminaHiSeq) was derived from the TCGA Research Network: http://cancergenome.nih.gov. The data set derived from this resource that supports the findings of this study is available at https://genome- cancer.ucsc.edu. All other data supporting the findings of this study are available from the corresponding author on reasonable request.
